# Prior experience with behavioral control over stress facilitates social dominance

**DOI:** 10.1101/2023.06.06.543982

**Authors:** Philip T. Coleman, Gabriel W. Costanza-Chavez, Heather N. Martin, Jose Amat, Matthew G. Frank, Rory J. Sanchez, Garrett J. Potter, Simone M. Mellert, Rene K. Carter, Gianni N. Bonnici, Steven F. Maier, Michael V. Baratta

## Abstract

Dominance status has extensive effects on physical and mental health, and an individual’s relative position can be shaped by experiential factors. A variety of considerations suggest that the experience of behavioral control over stressors should produce winning in dominance tests and that winning should blunt the impact of later stressors, as does prior control. To investigate the interplay between competitive success and stressor control, we first examined the impact of stressor controllability on subsequent performance in a warm spot competition test modified for rats. Prior experience of controllable, but not physically identical uncontrollable, stress increased later effortful behavior and occupation of the warm spot. Controllable stress subjects consistently ranked higher than did uncontrollable stress subjects. Pharmacological inactivation of the prelimbic (PL) cortex during behavioral control prevented later facilitation of dominance. Next, we explored whether repeated winning experiences produced later resistance against the typical sequelae of uncontrollable stress. To establish dominance status, triads of rats were given five sessions of warm spot competition. Reversible inactivation of the PL or NMDA receptor blockade in the dorsomedial striatum led to a long-term reduction in social rank. Stable dominance blunted the later stress-induced increase in dorsal raphe nucleus serotonergic activity, as well as prevented stress-induced social avoidance. In contrast, endocrine and neuroimmune responses to uncontrollable stress were unaffected, indicating a selective impact of prior dominance. Together, these data demonstrate that instrumental control over stress promotes later dominance, but also reveal that winning experiences buffer against the neural and behavioral outcomes of future adversity.

## Introduction

Dominance status has important consequences for an individual’s access to resources, health outcomes, and rates of reproductive success and survival (Sapolsky, 2005; Rivers and Josephs, 2010). There is considerable interest in the mechanisms by which experiential/behavioral variables influence hierarchy formation, as well as how dominance status impacts an individual’s response to future adversity. The medial prefrontal cortex (mPFC) has been implicated in (i) driving the effortful behavior necessary to attain a higher dominance rank in a social competition and (ii) the ability to generalize this rank to novel social contests (Cooper et al., 2015; Zhou et al., 2017; Padilla-Coreano et al., 2022). Winning experiences activate distinct circuits within the prelimbic cortex (PL) and lead to changes in the synaptic strength of layer V pyramidal neurons, the main output layer of the mPFC (Wang et al., 2011; Garcia-Font et al., 2022; Zhang et al., 2022). Moreover, manipulations that inhibit or activate the PL result in a downward or upward movement in social rank, respectively (Zhou et al., 2017).

In addition, repeated winning experiences can buffer against *social* stress outcomes (Karamihalev et al., 2020; LeClair et al., 2021). This is noted here because an experience with behavioral control over adverse events also activates PL layer V output (Baratta et al., 2009), and this PL output activation produces enduring protection against both future social (Amat et al., 2010) and nonsocial stressors (Amat et al., 2006). Controllable (escapable, ES), but not physically identical uncontrollable (yoked inescapable shock, IS), stressors increase the intrinsic excitability of layer V pyramidal neurons in the PL (Varela et al., 2012). Moreover, intra-PL blockade of NMDA receptor activity or inhibition of its downstream effector pathway (ERK/MAPK) prevent ES-induced resistance against the typical neurochemical and behavioral outcomes of later IS that occurs in a novel environment (Christianson et al., 2014). Thus, ES elicits a generalized resistance to adversity experienced in new environments with new task demands.

Given that an initial experience of behavioral control over stress produces long-lasting alterations to the PL, then it might be expected that ES would facilitate later dominance in a social competition that depends upon PL activation. Furthermore, any effect of ES on later dominance should be dependent on PL activation *during* the initial experience of control. We also hypothesize that these effects are restricted to male rats, as control in females is not protective across a variety of parameters and does not activate the same circuitry (Baratta et al., 2018, 2019; McNulty et al., 2022). Considering that dominance procedures like the tube test use food reinforcers during training, which can be devalued by stress, we examined how controllability impacts performance in a dominance task that does not involve appetitive motivation. We used a modified version of the warm spot test (WST) for rats, in which a triad competes for sole occupancy of a warm spot on a cold cage floor (Zhou et al., 2017).

We also investigated whether repeated winning in the warm spot competition produces similar resilience phenomena as does behavioral control. Specifically, we examined if the circuits involved in the enduring effects of ES are also engaged in the development of repeated winning. Additionally, we addressed whether stable winning buffers against the behavioral and neurochemical outcomes of IS. Robust activation of serotonin (5-HT) neurons in the dorsal raphe nucleus (DRN) is necessary and sufficient to elicit the behavioral sequelae of IS (for review, see Maier and Watkins, 2005), including IS-induced social avoidance, the behavioral endpoint measured here (Christianson et al., 2008). Thus, we measured extracellular levels of DRN 5-HT during IS and examined subsequent social interaction. The overarching hypothesis is that if behavioral control over *nonsocial* stressors (shock) and the experience of *social* winning activate the same prefrontal circuitry, they should be fungible.

## Materials and Methods

### Subjects

Adult male (275-300 g) and female (225-250 g) Sprague–Dawley rats (Envigo) were pair-housed on a 12-h light–dark cycle (lights on at 0700 h) with *ad libitum* access to standard laboratory chow and water. Only subjects implanted with guide cannula were individually housed to avoid potential damage from cage mates. Rats were allowed to acclimate to colony conditions for at least one week before experimentation. All behavioral procedures were carried out between 0900 and 1400 h. All experiments were in accordance with the National Institutes of Health *Guide for the Care and Use of Laboratory Animals* and were approved by the University of Colorado Boulder Institutional Animal Care and Use Committee.

### Wheel-turn ES/yoked IS procedure

For manipulation of stressor controllability, subjects were run in a triadic design as previously described (Baratta et al., 2007). One subject of each triad received ES, a second received yoked IS, and a third remained undisturbed in the colony room (home cage, HC). Each ES and IS rat were placed in a 14 × 11 × 17 cm (length × width × height) Plexiglas box with a wheel mounted in the front. The tail was secured to a Plexiglas rod extending from the back of the box and affixed with two copper electrodes and electrode paste. The single stress session consisted of 100 tailshock trials administered by a current-regulated shocker (Coulbourn Instruments). Tailshocks (1.0 mA) were presented on a 60-s variable interval schedule. Initially, the shock could be terminated by a quarter turn of the wheel by the ES rat. When trials were completed in less than 5 s, the response requirement was increased by one-quarter turn of the wheel, up to a maximum of four full turns of the wheel. The requirement was reduced if the trial was not completed in less than 5 s. If the trial was not completed in 30 s, the shock was automatically terminated, and the requirement was reset to a one-quarter turn of the wheel. For yoked IS rats, the onset and offset of each tailshock were identical to those of its ES partner. A computer equipped with Graphic State 4 (Coulbourn Instruments) controlled the experimental events and recorded the wheel turn requirement and escape latency for each trial.

### Inescapable shock procedure

For behavioral experiments that only involved inescapable stress, rats were placed in a Plexiglas restraint tube (17.5 cm in length × 6.0 cm in diameter) with a Plexiglas rod protruding from the rear to which the rat’s tail was taped and affixed with two copper electrodes. Rats received a single session of 100 inescapable tailshocks (5 s duration, 1.0 mA each) with a variable inter-trial interval ranging from 30–90 s (average of 60 s).

### Warm spot

The warm spot competition was adapted from the protocol previously described in mice (Zhou et al., 2017). An empty housing cage (48.3 cm in length × 26.7 cm in width × 20.3 cm in height) was placed on ice to cool the cage floor (0°C). In one corner of the cage, an inverted circular lid (diameter: 8.9 cm; peripheral lip height: 1.6 cm) containing a thin toe warmer covered with paper nesting material served as the ‘warm spot’ (∼35°C). The warm spot was sufficiently large to accommodate only a single adult rat and was affixed to the cage floor to prevent displacement during the competition. Prior to experimentation, rats were individually habituated to the above warm spot-cold cage environment for 20 min.

For all experiments, the warm spot competition involved a triad of non-cage-mates. Triads were first introduced into an empty housing cage with a cold floor that did not contain a warm spot. After 10 min, the triad was transferred to the warm spot-cold cage environment in which subjects competed over a 20 min period for access to the warm spot. All sessions were videotaped to minimize experimenter presence in the procedure room. Subjects were marked with different colors for unique identification within the triad and to ensure videos were scored blinded to treatment. Occupancy time of the spot for each triad subject was calculated and expressed as a percentage of the total time the warm spot was occupied during the 20 min session. Competition behavioral measures included the number of 1) pushes initiated (not in response to being pushed by another triad member); 2) resistance bouts (either withstanding or pushing back after being pushed); and 3) retreats (withdrawing from warm spot in response to the actions of another triad member).

### Juvenile social exploration (JSE)

As described in Christianson et al. (2010), animals were moved from their colony room to a novel procedure room (150 lux at the position of the animal) and placed in a standard Plexiglas housing cage with bedding and a wire lid. Following a 1 h habituation period, a juvenile stimulus rat (28-35 days old male Sprague-Dawley) was introduced to the cage for 3 min and exploratory behaviors (sniffing, pinning, chasing, and allogrooming) initiated by the adult were scored by an observer blind to treatment.

### Stereotactic surgery

All surgeries were performed under inhaled isoflurane anesthesia (5% induction, 2% maintenance in 2.5 L/min O_2_; Piramal Critical Care). For drug microinfusion studies, cannula (26 gauge; P1 Technologies) were implanted bilaterally in either the PL (A/P: +2.6; M/L: +/-0.5; D/V: -1.8 mm from the pial surface) or DMS (A/P: -0.2; M/L: +/-2.1; D/V: -3.0 mm from the pial surface) and secured to the skull with stainless steel screws and acrylic cement. Internal guide cannulae were inserted to keep the cannula patent and were held in place with a fitted dust cap (P1 Technologies). Subjects were given 10-14 days to recover before experimentation. For *in vivo* microdialysis, general surgical procedures for guide cannula implantation into the midline DRN (A/P: -7.8, M/L: 0.0, D/V: -4.9 mm from the pial surface) were similar to above. However, a screw cap from a 15-mL conical centrifuge tube (with the central portion removed) was affixed to the skull in an inverted orientation so that it encircled the guide cannula. This was done to protect the microdialysis guide cannula during tailshock. Subjects were allowed 7-10 days to recover. All subjects were given postoperative subcutaneous injections of an extended-release nonsteroidal anti-inflammatory (meloxicam, 4.0 mg/kg; Vetmedica) and an antibiotic (CombiPen-48, 0.25 mL/kg; Bimeda). At the end of the experiment, brains were collected, sliced at 30 µm, and stained with cresyl violet for verification of cannula placement. Subjects were only included in the data analysis if tissue damage from the cannula tip fell within the target structure.

### Drug microinfusion

For pharmacology studies, subjects received microinjections 30-45 min prior to stress treatment or warm spot competition. Microinjections were made in a quiet room near the testing area. Subjects were gently restrained and microinjectors (33 gauge; P1 Technologies), attached to a 25-μl Hamilton syringe with PE 50 tubing, that extended 1 mm beyond cannulae tips were inserted. Syringes were mounted in a Kopf microinjection unit (Model 5000) and delivered either the GABA_A_ agonist muscimol (50 ng/0.5 μl/side; Sigma-Aldrich) to the PL, the NMDA antagonist AP5 (3 μg/0.5 μl/side; Tocris Bioscience) to the DMS, or equal volume sterile saline to either brain region. Injectors were left in place for 80 s following injection to allow for diffusion. All drugs were dissolved in 0.9% sterile saline and doses were chosen based on our prior work in male Sprague-Dawley rat (Amat et al., 2005, 2014).

### In vivo microdialysis

A CMA 12 microdialysis probe (0.5 mm diameter, 1 mm length, 20 kDa cut-off) was inserted through the cannula guide to the midline of the DRN the afternoon before sample collection. A portion of a 15-mL conical tube was screwed onto the skull-mounted screw cap, through which the dialysis tubing, protected within a metal spring, was entered and attached to the probe. Each subject was placed individually in a Plexiglas bowl and infused with artificial cerebrospinal fluid (pH 7.2, 145.0 mM NaCl, 2.7 mM KCl, 1.2 mM CaCl, 1.0 mM MgCl_2_) at a rate of 0.2 μL min^-1^. The following day the flow rate was increased to 0.8 μL min^-1^. After a 90-min stabilization period, four baseline samples were collected. Subjects were then transferred to a Plexiglas open-top box where they received 100 trials of inescapable tailshocks (5 s duration, 1.0 mA, 60 s average inter-trial interval) or no stress. Following stress treatment, subjects were moved back to the bowls, where three additional samples were collected. During the baseline, stress, and post-stress phases, dialysates were collected at 20 min intervals and stored in an -80°C freezer until analysis. Microdialysis data are expressed as a percentage of baseline, defined as the mean of four consecutive samples collected prior to the stress phase.

### 5-HT analysis

5-HT concentration was measured in dialysates using high-performance liquid chromatography (HPLC) with electrochemical detection. The system consisted of an online Shimadzu DGU-2045 degasser, an ESA 584 pump, a Dionex UltiMate 3000 electrochemical detector with a 6041 RS ultra amperometric cell and autosampler, and an ESA 5020 guard cell. The analytical column was an Acclaim RSLC PolarAdvantage II (2.1 × 100 mm; Thermo Fisher Scientific), maintained at 38°C, and the mobile phase was the ESA buffer MD-TM. The analytical cell potential was kept at +220 mV, and the guard cell at +250 mV. External standards (Sigma-Aldrich) dissolved in artificial cerebrospinal fluid were run each day to quantify 5-HT.

### Tissue dissection

Rats were given a lethal dose of sodium pentobarbital. Cardiac blood was collected prior to performing transcardial perfusion with ice-cold saline (0.9%) for 3 min to remove peripheral immune leukocytes from central nervous system vasculature. Brain was rapidly extracted, placed on ice, and the hypothalamus dissected. Tissue samples were flash frozen in liquid nitrogen and stored at -80°C.

### Enzyme-linked immunosorbent assay (ELISA)

Cardiac blood was centrifuged (10 min, 14,000 × g, 4°C) and serum stored at -80°C. Serum corticosterone (CORT) was measured using a competitive immunoassay (Enzo Life Sciences) as described in the manufacturer’s protocol. Serum CORT levels were expressed as μg/dL.

### Quantitative real-time PCR

Total RNA was isolated from hypothalamus using TRI Reagent (MilliporeSigma) and a standard method of phenol:chloroform extraction (Chomczynski and Sacchi, 1987). Total RNA was quantified using a NanoDrop 2000 spectrophotometer (Thermo Fisher Scientific). cDNA synthesis was performed using the SuperScript II Reverse Transcriptase kit (Thermo Fisher Scientific). A detailed description of the PCR amplification protocol has been published previously (Frank et al., 2006). cDNA sequences were obtained from Genbank at the National Center for Biotechnology Information (NCBI; www.ncbi.nlm.nih.gov). Primer sequences were designed using the Operon Oligo Analysis Tool (http://www.operon.com/tools/oligo-analysis-tool.aspx) and tested for sequence specificity using the Basic Local Alignment Search Tool at NCBI (Altschul et al., 1997). Primers were obtained from Thermo Fisher Scientific, and specificity was verified by melt curve analyses. All primers were designed to span exon/exon boundaries and thus exclude amplification of genomic DNA. Primer sequences: interleukin-1β (*Il1b*), F: CCTTGTGCAAGTGTCTGAAG, R: GGGCTTGGAAGCAATCCTTA; interleukin-6 (*Il6*), F: AGAAAAGAGTTGTGCAATGGCA, R: GGCAAATTTCCTGGTTATATCC; C-X3-C motif chemokine receptor 1 (*cx3cr1*), F: AGCTGCTCAGGACCTCACCAT, R: CCGAACGTGAAGACAAGGGAG; β-actin (*Actb)*, F: TTCCTTCCTGGGTATGGAAT, R: GAGGAGCAATGATCTTGATC. PCR amplification of cDNA was performed using the Quantitect SYBR Green PCR Kit (Qiagen). Formation of PCR products was monitored in real time using the CFX96 Touch Real-Time PCR Detection System (BioRad). Relative gene expression was determined using *Actb* as the housekeeping gene and the 2^-ΔΔCT^ method (Livak and Schmittgen, 2001).

### Statistical analysis

Data analyses were performed using Prism software (GraphPad, RRID:SCR_002798). The effect of treatment was analyzed with unpaired *t* test, one-way, two-way, or repeated measures ANOVA. Main effects and interactions were considered significant if *p* < 0.05. When appropriate, *post-hoc* analyses were performed with Tukey’s multiple comparison test or unpaired *t* test. For nonparametric data (latency to spot, rank), Mann-Whitney U or Kruskal-Wallis (for comparison of more than two groups) followed by Dunn’s multiple comparison test were used. In all cases, data are expressed as the mean ± SEM.

## Results

### Effects of stressor controllability on later social dominance

To evaluate the impact of stressor controllability on subsequent dominance behavior, male rats were individually habituated to the warm spot apparatus 24 h prior to receiving ES, yoked-IS, or HC treatment (Fig. 1*A*). One week later ES/IS/HC triads were exposed to the warm spot competition. One-way ANOVA showed a significant effect of prior stress condition (*F*_2,30_ = 5.79, *p* = 0.007, *n* = 11 per group), with ES subjects spending significantly more time on the warm spot than both IS and HC subjects (*p*’s < 0.05; Fig. 1*B*). The impact of the stressor on later dominance was selective to controllable stress; the percentage of warm spot occupation did not differ between IS and HC subjects (*p* > 0.999). Importantly, once the competition was initiated, the latency time to first access the warm spot did not differ between groups, indicating that all groups exhibited similar levels of recall for the warm spot location (*p* = 0.120, Kruskal-Wallis; data not shown). Across triads, ES subjects consistently ranked higher than IS subjects (*p* = 0.014, Dunn’s; Fig. 1*C*) and exhibited a distinct behavioral profile during the 20 min WST competition. Prior ES increased the number of resistance behaviors (push-backs, maintaining spot occupation when pushed by another member of the triad) compared to IS and HC (*p*’s = 0.033; Fig. 1*D*). In contrast, there were no significant group differences in the mean number of pushes initiated (*F*_2,30_ = 0.883, *p* = 0.424) and passive responses (retreats from the warm spot; *F*_2,30_ = 1.179, *p* = 0.322; Fig. 1*D*).

**Figure 1.**
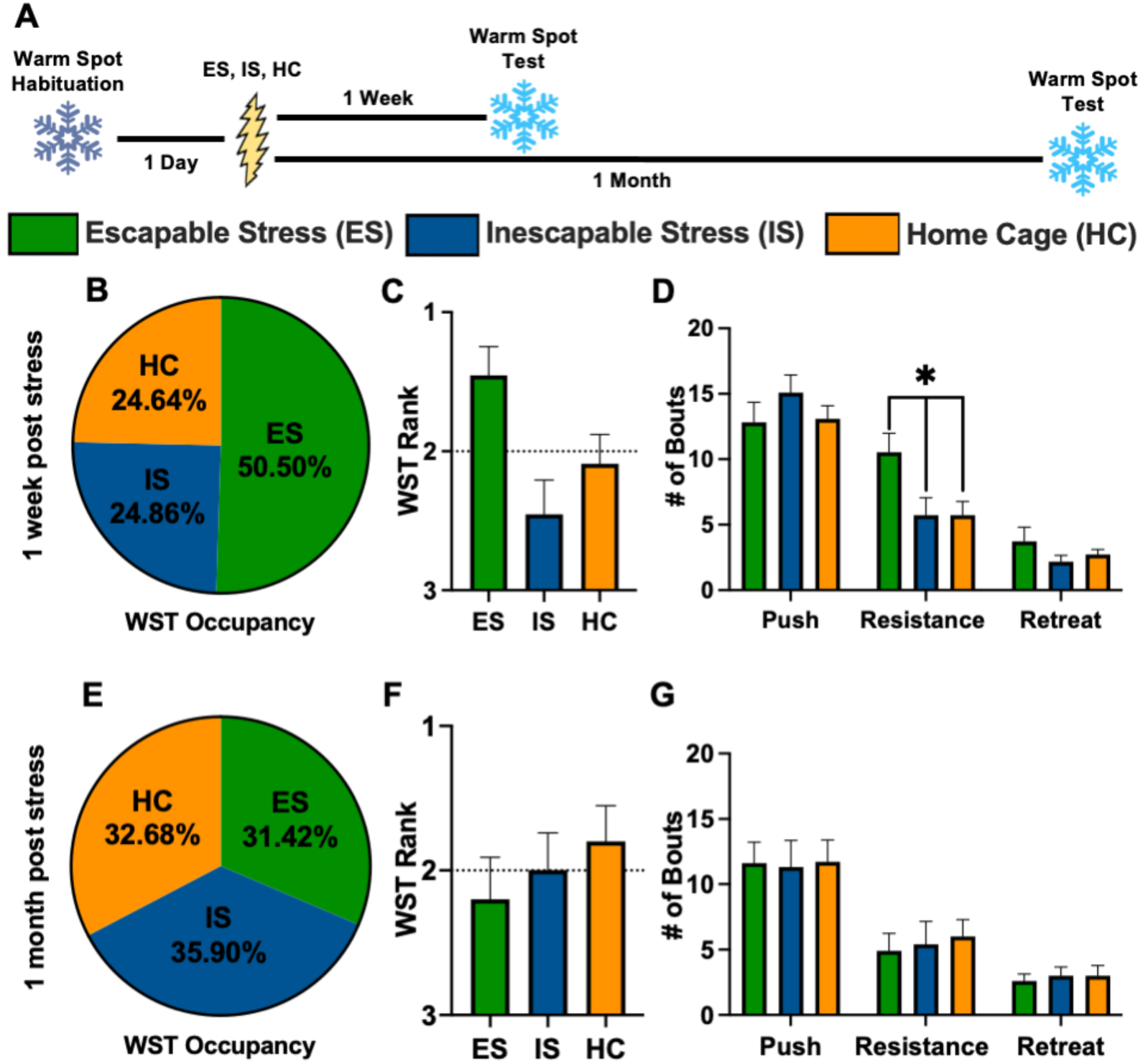
Behavioral control increases dominance in the warm spot competition 1 week later. ***A***, Experimental timeline. ***B***, Percent occupancy in the warm spot test (WST) 1 week after stress treatment. Each triad consisted of one member from escapable stress (ES), yoked-inescapable stress (IS), and no stress (home cage, HC) groups. ***C***, Average rank positions across warm spot triads. The dotted line indicates expected ranking if there was no impact of prior stress treatment. ***D***, Number of pushes initiated, resistance bouts (push-backs, withstanding a push) and retreats. ***E***, Percent occupancy in the WST 1 month after stress treatment. ***F***, Average rank positions across warm spot triads. ***G***, Number of individual behaviors. Values represent the mean ± SEM. \**p* < 0.05, Dunn’s (rank) and Tukey’s *post hoc* tests.

The ES-induced facilitation of dominance was both time-limited and sex-specific. In a separate male cohort, no significant group differences emerged when the warm spot competition occurred 30 days post-stress treatment (*n* = 10 per group; Fig. 1*E-G*). Warm spot occupation was similar between ES, yoked-IS, and HC groups (*F*_2,27_ = 0.085, *p* = 0.919). An impact of behavioral control phenomena is often reported to be absent in female rats (Fallon et al., 2020), and here we tested whether ES in females would produce dominance in the warm spot competition one week later as it does in males (Fig. 2*A*). Males and females typically perform the wheel-turn escape response with equal proficiency, and this held true here as well. Repeated-measures ANOVA indicated no differences in ES performance between females and males (male data from Fig. 1*B*) in the wheel-turn response requirement throughout the entire 100-trial session (sex: *F*_1,22_ = 0.008, *p* = 0.931; Fig. 2*B*). Despite a similar level of wheel-turn acquisition and motivation to escape across trials, prior ES in females had no impact on overall warm spot occupancy (*F*_2,39_ = 0.875, *p* = 0.425, *n* = 14 per group; Fig. 2*C*), rank (*p* = 0.390, Kruskal-Wallis; Fig. 2*D*), or individual behavioral measures (*p*’s > 0.05; Fig. 2*E*).

**Figure 2.**
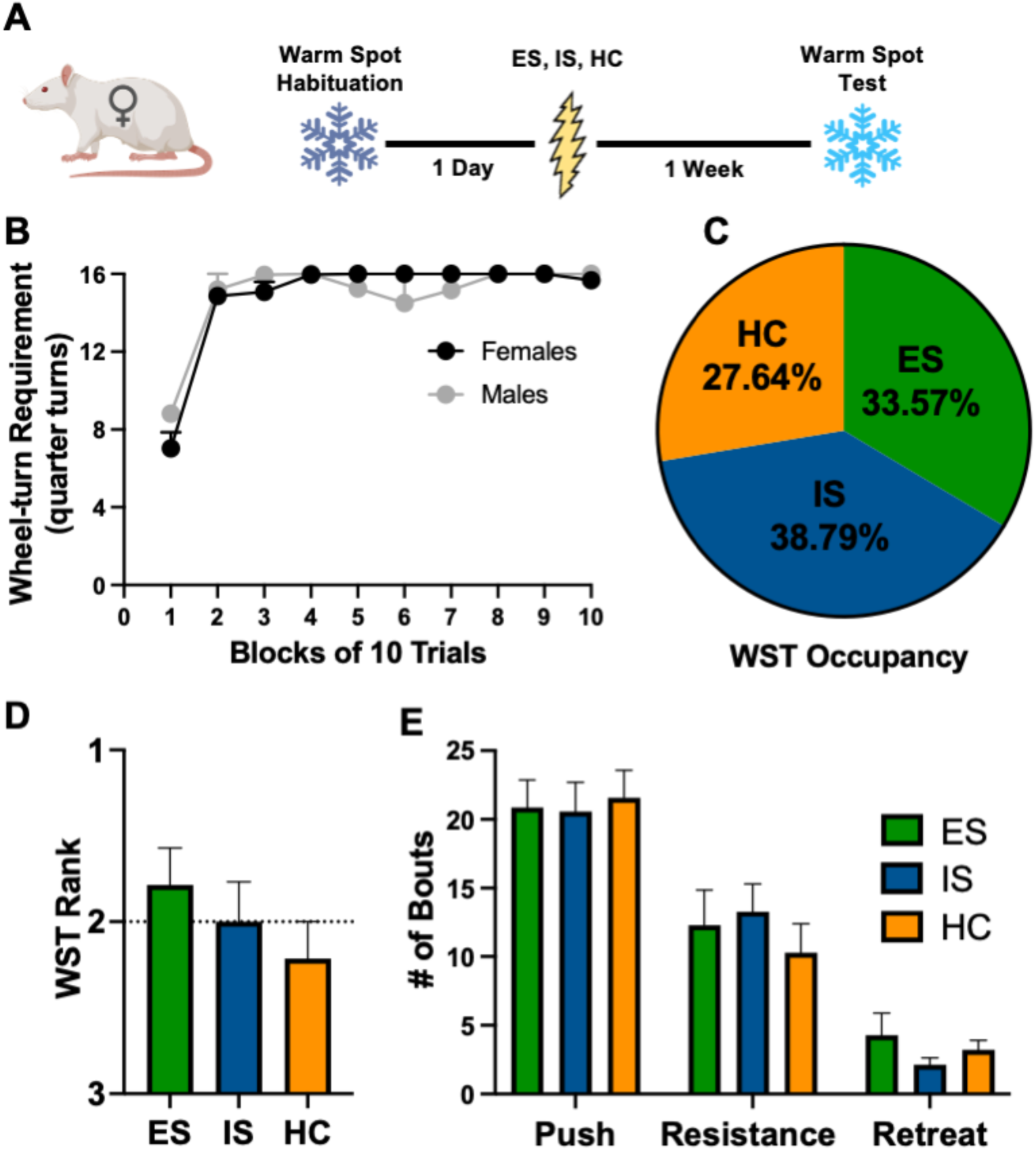
Behavioral control in female rats does not produce dominance in the warm spot competition. ***A***, Experimental timeline. ***B***, Comparison of female wheel-turn escape behavior with that of males in Figure 1. Number of quarter turns of the wheel attained as the escape requirement for each trial. ***C***, Percent occupancy of the warm spot test (WST) for escapable stress (ES), yoked-inescapable stress (IS), and no stress (home cage, HC) groups 1 week after stress treatment. ***D***, Average rank positions across warm spot triads. The dotted line indicates expected ranking if there was no impact of prior stress treatment. ***E***, Number of individual behaviors. Values represent the mean ± SEM.

### Prelimbic cortex involvement in the dominance-producing effects of behavioral control

A variety of data supports the idea that the enduring effects of behavioral control requires PL activity during the control experience (Baratta et al., 2009; Christianson et al., 2014). Thus, we addressed whether pharmacological inactivation of the PL during ES would prevent its selective impact on later warm spot behavior (Fig. 3*A*). ES subjects received intra-PL bilateral microinfusions of either muscimol (ES-M; 50 ng/0.5 μl/side, GABA_A_ agonist) or saline vehicle (ES-V) prior to stress (cannula placements shown in Fig. 3*B*). IS and HC triad members received only saline vehicle. Importantly, ES subjects with PL inactivation learned the escape (control) response quite well. Efficiency in wheel-turn escape behavior was identical between ES-V and ES-M groups (drug: *F*_1,17_ = 0.332, *p* = 0.572; Fig. 3*C*). Once again, ES-V produced dominance 1 week later in the 20-min WST compared to IS-V and HC-V groups (*F*_2,30_ = 8.261, *p* = 0.001, *n* = 11 per group; Fig. 3*D*). ES-V spent a significantly greater amount of time on the warm spot compared to HC-V (*p* = 0.035) and IS-V (*p* = 0.001). In contrast, intra-PL muscimol given before ES eliminated the dominance-producing effect of ES; that is, ES subjects no longer showed increased spot occupation times (Fig. 3*E*). These conclusions were confirmed by ANOVA (*F*_2,21_ = 12.52, *p* < 0.001, *n* = 8 per group). HC-V now had significantly greater occupancy time than ES-M and IS-V groups (*p* = 0.005 and *p* < 0.001, respectively), whereas ES-M and IS-V did not differ (*p* = 0.456). Intra-PL muscimol also reduced the effects of ES on later rank (*p* = 0.029, Mann-Whitney U test; Fig. 3*F*) and the number of resistance bouts (*p* = 0.036, unpaired *t* test; Fig. 3*G*).

**Figure 3.**
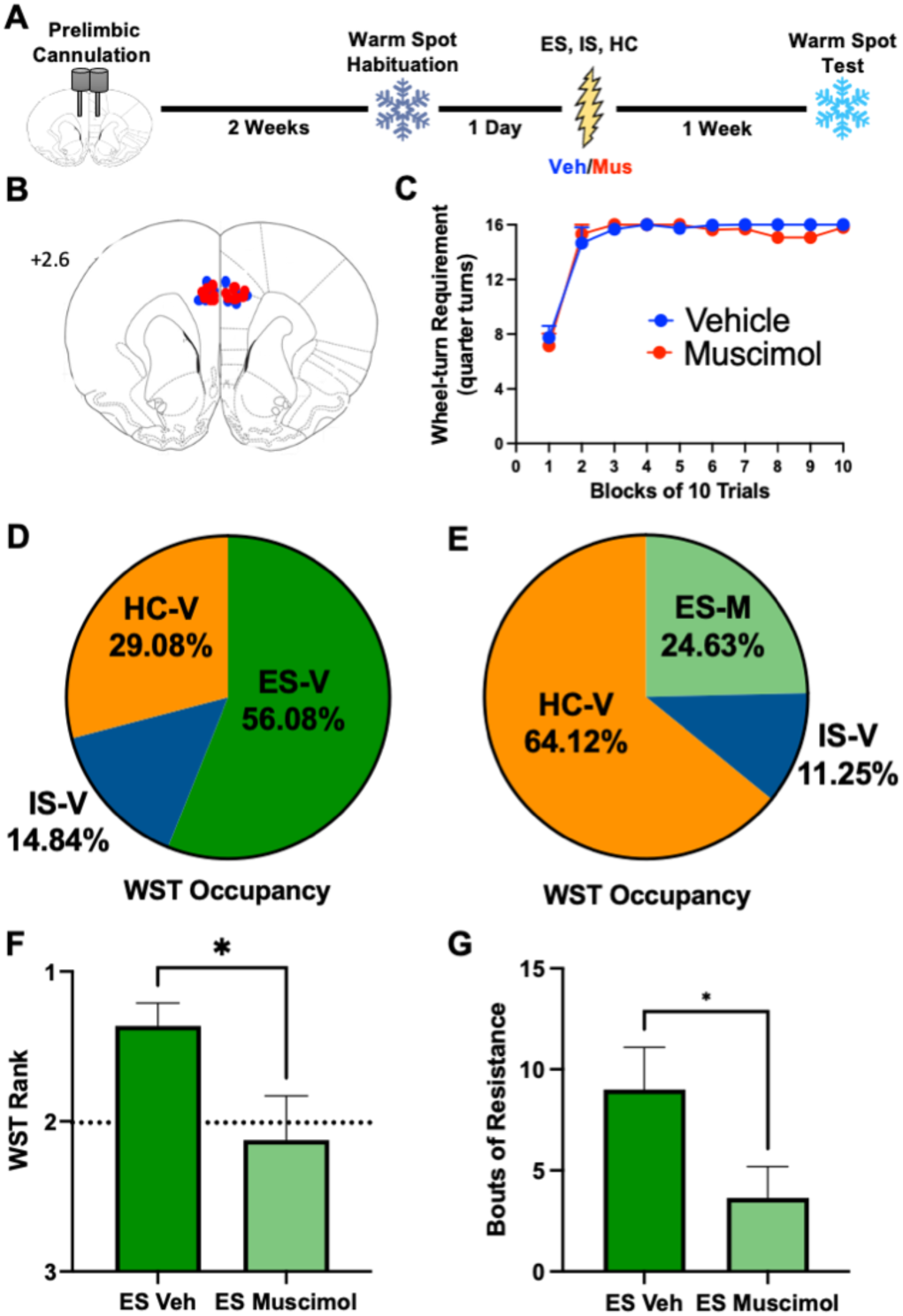
Prefrontal involvement in the dominance-producing effects of behavioral control. ***A***, Experimental timeline. ***B***, Cannula placement in the prelimbic region of the medial prefrontal cortex for vehicle- (blue dots) and muscimol-treated (red dots) escapable stress (ES) subjects. Number indicates distance (mm) anterior to bregma. ***C***, Comparison of wheel-turn escape behavior between intra-prelimbic vehicle- and muscimol-treated ES subjects. Number of quarter turns of the wheel attained as the escape requirement for each trial. ***D***, Percent occupancy of the warm spot test for triads in which the ES subject received vehicle or ***E***, muscimol. ***F***, Average rank position of ES vehicle and muscimol groups. The dotted line indicates expected ranking if there was no impact of prior stress treatment. ***G***, Number of resistance bouts. Values represent the mean ± SEM. \**p* < 0.05, Mann-Whitney U (rank) and unpaired *t* test (resistance).

### Stable dominance in the warm spot requires the corticostriatal system

Similar to an experience with behavioral control, prior history of winning in a social competition can also increase the probability of winning in future contests (’winner effect’; Landau, 1951; Dugatkin, 1997). We next addressed whether corticostriatal structures are involved in the development of sustained winning in the warm spot test since both the PL and DMS are necessary for the protective effects of behavioral control (Amat et al., 2014; Christianson et al., 2014). First, rats were implanted with bilateral cannula in the PL (Fig. 4*B*) 2 weeks prior to a series of 20-min warm spot triad competitions. There was a total of 5 competitions, each separated by 48 h. Winners in the initial warm spot session (subjects that had the highest occupancy time in each triad) received either intra-PL muscimol or vehicle 30 min prior to the second session. All other members of the triad received saline vehicle. In session 1 winners, vehicle treatment had no impact on subsequent performance in session 2. PL inactivation, however, produced a marked decrease in the percentage of spot occupancy. Repeated-measures ANOVA indicated a significant main effect of drug condition (*F*_1,13_ = 7.993, *p* = 0.014, *n* = 7-8 per group), but no significant interaction between drug and session (*F*_4,52_ = 1.819, *p* = 0.139; Fig. 4*D*). Surprisingly, muscimol treatment not only reduced session 2 dominance (*p* = 0.005; Fig. 4*E*), but also in subsequent sessions when subjects were drug-free (*p* = 0.017; Fig. 4*F*).

**Figure 4.**
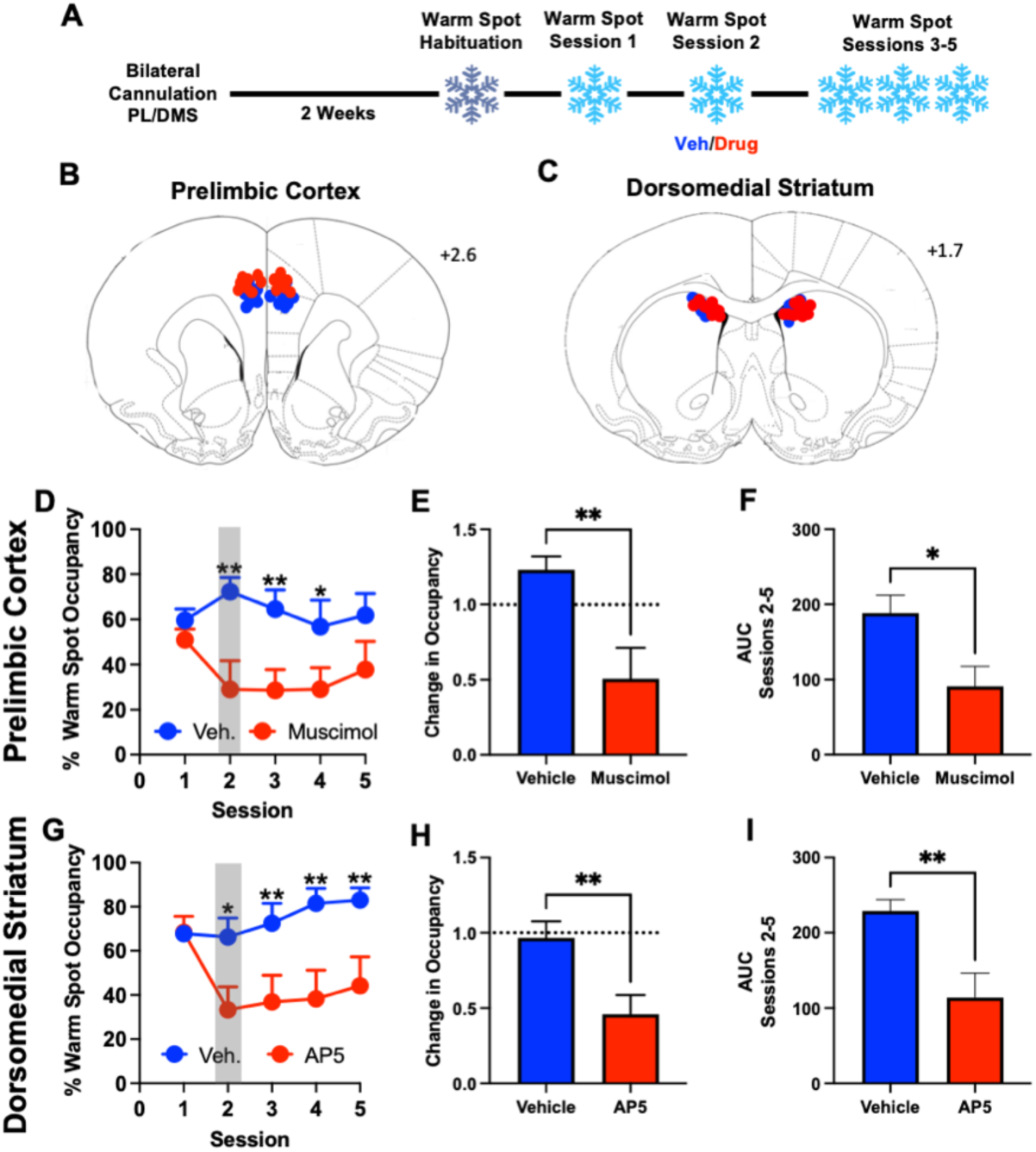
The prelimbic cortex and the dorsomedial striatum are required for the development of stable dominance. ***A***, Experimental timeline. ***B***, Cannula placements for session 1 winners that received vehicle (blue dots) or muscimol (red dots) in the prelimbic cortex (PL) cortex or ***C***, vehicle (blue dots) or AP5 (red dots) in the dorsomedial striatum (DMS). Numbers indicate distance (mm) anterior to bregma. ***D***, Percent occupancy of the warm spot test for initial winners that received intra-PL vehicle or muscimol prior to session 2. Gray bar indicates drug-treatment session. ***E***, Change in spot occupancy (session2/session1) for initial session winners. ***F***, Histogram depicting the mean area under the curve (AUC) for warm spot occupancy, sessions 2-5 only. ***G***, Percent occupancy of the warm spot test for initial winners that received intra-DMS vehicle or muscimol prior to session 2. Gray bar indicates drug-treatment session. ***H***, Change in spot occupancy (session2/session1) for initial session winners. ***I***, Histogram depicting AUC for warm spot occupancy, sessions 2-5. Values represent the mean ± SEM. \**p* < 0.05, \*\**p* < 0.01, unpaired *t* test.

Similar to the above, a second cohort was implanted with bilateral cannula targeting the DMS rather than the PL (Fig. 4*C*). Half of the winners from the first warm spot session received intra-DMS microinjections of AP5 prior to session 2, whereas the remaining half received saline vehicle. Repeated-measures ANOVA showed significant effects of drug condition (*F*_1,13_ = 10.26, *p* = 0.007, *n* = 7-8 per group) and interaction between drug and session (*F*_4,52_ = 3.137, *p* = 0.022; Fig. 4*G*). Once again, vehicle-treated session 1 winners continued to show high levels of spot occupancy across all sessions. Intra-DMS AP5 decreased occupation time during session 2 (*p* = 0.009; Fig. 4*H*), and this loss of dominance was evident throughout the remaining sessions (*p* = 0.005; Fig. 4*I*).

### Impact of repeated winning on the behavioral and neurochemical sequelae of inescapable stress

Given the involvement of the PL and DMS in the development of stable dominance, and the prior work showing the necessity of these structures for the occurrence of the protective effects of behavioral control, we next sought to determine if the experience of repeated winning buffers against the behavioral and neurochemical outcomes of future IS, as does prior ES (Fig. 5*A,B*). Triads received 5 daily warm spot competitions prior to receiving a single session of IS (100 trials, 1.0 mA and 5 s duration each) or no stress (HC). JSE was assessed 24 h later. As is typical, exposure to IS reduced JSE in rats that did not undergo the warm spot procedure (IS-No WST; Fig. 5*C*). Prior history of repeated winning (top rank position for at least 4 out of the 5 sessions) completely blocked the social avoidance produced by IS. Importantly, the impact was specific to winning. Repeated losing (lowest rank position for at least 4 out of the 5 sessions) led to reduced social interaction, independent of stress treatment (data not shown). To control for the reduced amount of time spent on the cold floor by winners, we also included a group in which subjects were matched to stable winners such that they received an equivalent amount of time exposed to both the warm spot and cold cage floor but did not have to compete for gaining spot access (“Exposure”). Matching exposure to the warm spot-cold cage environment to that of winners did not buffer against IS-induced social avoidance. These conclusions were confirmed by ANOVA showing a main effect of stress condition (*F*_1,47_ = 10.35, *p* = 0.002, *n* = 8-10 per group) and a significant interaction between stress and warm spot experience (*F*_2,47_ = 4.351, *p* = 0.02; Fig. 5*C*).

**Figure 5.**
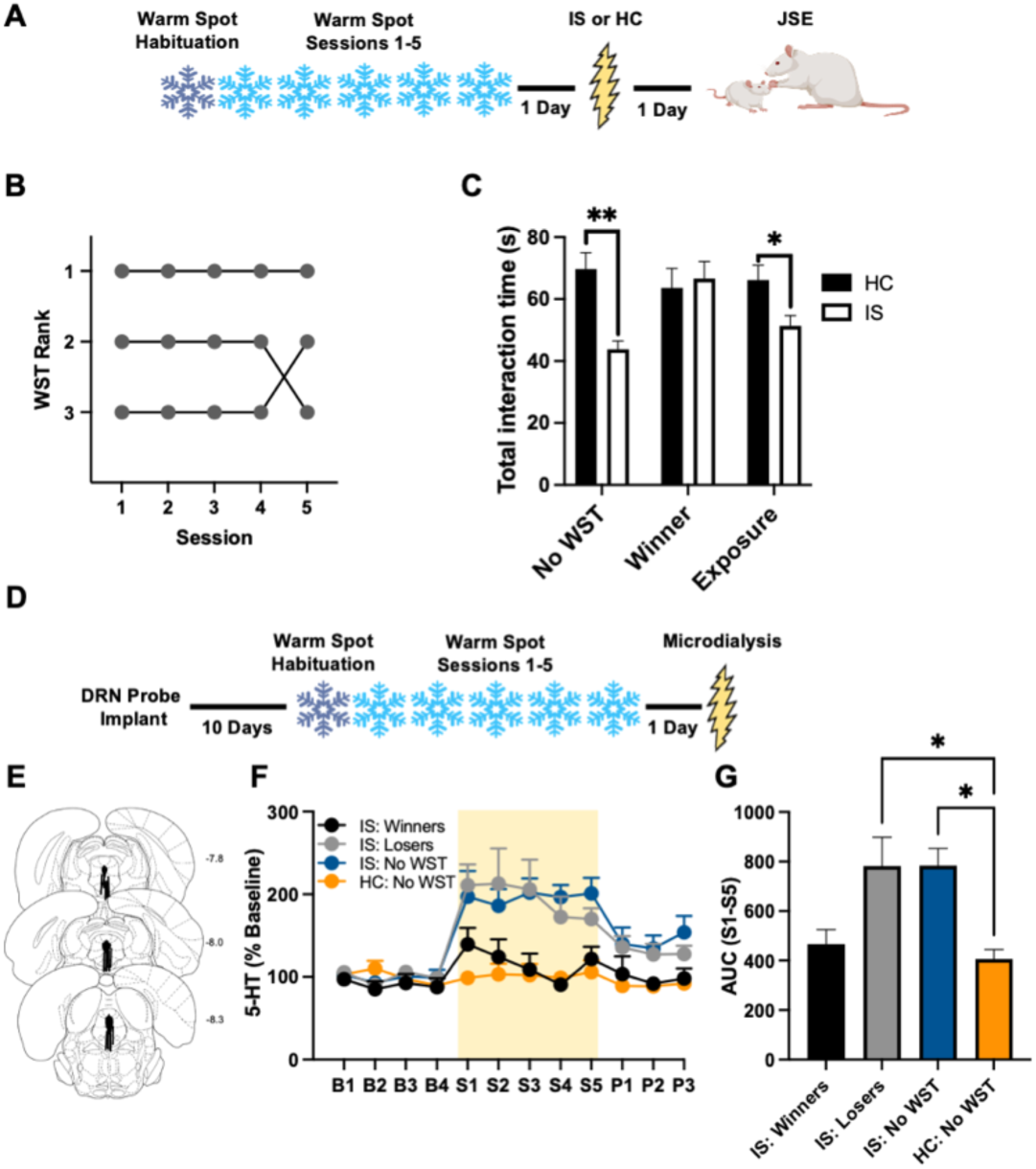
Stable dominance buffers against the behavioral and neurochemical outcomes of inescapable shock. ***A***, Experimental timeline of behavioral experiment. Warm spot competition “naïve” subjects (No WST), repeated winners, repeated losers, and controls that received equal exposure to the warm spot environment but not competition were given inescapable stress (IS) or no stress (home cage, HC). ***B,*** Representative rankings across 5 sessions for triads with a repeated winner (left) and a repeated loser (right) across five sessions. ***C***, A 3-min juvenile social exploration (JSE) session was conducted 24 h after stress treatment and data is expressed as the total exploration time. ***D***, Experimental timeline of *in vivo* microdialysis experiment. ***E***, Black bars represent the placement of microdialysis probes in the dorsal raphe nucleus (DRN). Numbers indicate distance (mm) posterior to bregma. ***F***, Extracellular serotonin (5-HT) as a percentage of baseline in the mid-caudal region of the DRN during IS or HC. Samples were collected every 20 min during the baseline (B1-B4), stress (S1-S5; depicted by yellow shaded area), and post-stress (P1-P3) phases. ***G***, Histogram depicting the mean area under the curve (AUC) for extracellular 5-HT during IS (S1– S5). Values represent the mean ± SEM. \**p* < 0.05, \*\**p* < 0.01, Tukey’s.

Exposure to IS produces a persistent increase in DRN 5-HT efflux indicative of DRN 5-HT activation, and activation of DRN 5-HT neurons is both necessary and sufficient for the behavioral sequelae of IS, such as social avoidance (Maswood et al., 1998; Christianson et al., 2008; Baratta et al., 2023). To examine whether a history of winning also buffers against IS-induction of DRN 5-HT efflux, subjects were first implanted with a microdialysis probe targeted to the mid-caudal region of the DRN (Fig. 5*E*). IS led to a large increase in extracellular DRN 5-HT that remained elevated throughout the entire stress treatment (S1-S5; group: *F*_3,23_ = 5.358, *p* = 0.006, *n* = 5-9 per group; Fig. 5*F*), however this only occurred in repeated warm spot losers (IS-Losers) and subjects that did not undergo warm spot competition (IS-No WST). In contrast, IS led to an initial increase in 5-HT that quickly returned to baseline in subjects with a history of winning (IS-Winners). Comparison of the mean area under the curve (*F*_3,23_ = 5.205, *p* = 0.007; Fig. 5*G*) further showed that IS only elevated 5-HT levels in IS-Losers and IS-No WST groups (*p*’s < 0.05), whereas IS-Winners did not differ from no stress subjects (HC-No WST; *p* = 0.987).

### Effect of stable dominance on the glucocorticoid and neuroimmune responses to inescapable stress

In addition to DRN 5-HT activity, exposure to IS also induces an array of endocrine (Maier et al., 1986; Fleshner et al., 1995) and central immune responses (O’Connor et al., 2003; Frank et al., 2019). Here we addressed whether the stress-buffering effects of stable dominance (again, top rank position for at least 4 out of the 5 sessions) extends to IS-induced endocrine and neuroimmune changes that are known not to be buffered by behavioral control. Immediately after the IS procedure (or no stress), cardiac blood was collected to obtain a measure of circulating CORT, along with tissue dissection of the hypothalamus to examine cytokine (IL-1β, IL-6) and chemokine (CX3CR1) mRNA expression. As is typical, IS led to a pronounced increase of serum CORT (*F*_1,27_ = 541.8, *p* < 0.001, *n* = 7-9 per group), but there was no main effect of prior dominance status (*F*_1,27_ = 1.685, *p* = 0.205) nor an interaction between status and stress conditions (*F*_1,27_ = 2.207, *p* = 0.149; Fig. 6*B*). The elevation of CORT elicited by IS was similar in both stable winners and losers. IS also increased IL-1β (*F*_1,26_ = 36.02, *p* < 0.001, *n* = 7-8 per group), but not IL-6 (*F*_1,26_ = 4.178, *p* = 0.051), mRNA levels in the hypothalamus (Fig. 6*C*). Once again, the stress-induced increase in hypothalamic IL-1β mRNA was independent of prior winning history (*F*_1,26_ = 0.029, *p* = 0.867) nor was there an interaction between status and stress (*F*_1,26_ = 0.166, *p* = 0.687). We also measured the gene expression level of the chemokine receptor CX3CR1. Expressed on the surface of microglia, the primary innate immune cell of the brain, CX3CR1 is thought to constitutively maintain microglia in a quiescent state through interactions with its ligand expressed on neurons (Cardona et al., 2006). IS-induced neuroinflammation may be mediated, in part, by IS-induced downregulation of CX3CR1. IS significantly downregulated CX3CR1 gene expression (*F*_1,26_ = 59.78, *p* < 0.001; Fig. 6*C*) and this reduction did not differ between winners and losers.

**Figure 6.**
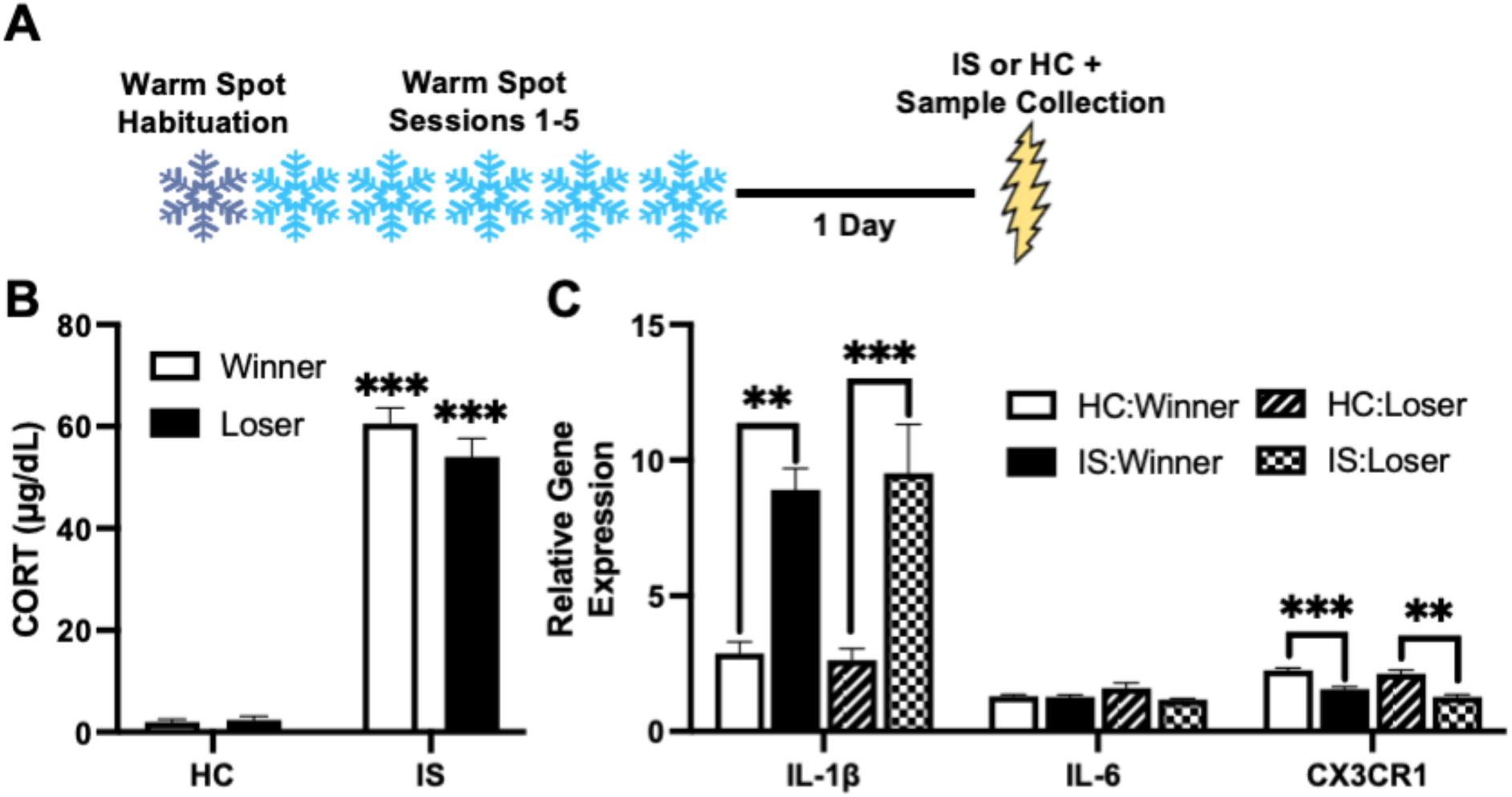
Dominance status does not impact endocrine and neuroinflammatory sequelae of inescapable shock. ***A***, Experimental timeline. Repeated winners and losers in the warm spot test (WST) were given inescapable stress (IS) or no stress (home cage, HC). ***B***, Measurement of serum corticosterone (μg/dL, CORT) collected immediately after stress treatment. ***C***, Expression of neuroinflammatory-related genes in hypothalamus dissected immediately after stress treatment. Relative gene expression of (left to right) interleukin-1β (*Il1b*), interleukin-6 (*Il6*), and C-X3-C motif chemokine receptor 1 (*cx3cr1*) mRNA. Values represent the mean ± SEM. \*\**p* < 0.01, \*\*\**p* < 0.001 versus HC, Tukey’s.

## Discussion

The current set of experiments led to two primary findings. First, behavioral control in males increases the number of effortful behavioral epochs during future competitive interactions and facilitates their winning, an outcome that requires PL activation during the control experience. Second, the development of stable dominance involves corticostriatal structures (both PL and DMS), and once established, buffers against the typical neurochemical and behavioral sequelae of IS. We also observed that IS-induced endocrine and neuroimmune outcomes were unaffected by dominance rank, suggesting boundary conditions to the stress buffering effects of stable dominance.

Exposure to adverse events can shape future dominance hierarchies (Park et al., 2018; Šabanović et al., 2020). Our focus on the controllability of the stressor was motivated by the fact that a single experience of behavioral control leads to long-lasting resistance against future adversity in a prefrontal-dependent manner (Maier, 2015). Similarly, prefrontal circuits have been implicated in a number of features related to dominance behavior, such as competitive success and its transitivity (Wang et al., 2011; Zhou et al., 2017), effort expenditure (Zhou et al., 2017; Porter and Hillman, 2021), detection of the relative social rank of self and others (Garcia-Font et al., 2022; Padilla-Coreano et al., 2022), and working memory function that may contribute to these processes (D’Esposito et al., 2000; Miller, 2013). If behavioral control and winning strengthen and are regulated by the same circuits, then experiencing control should facilitate later winning. Indeed, we found that in males ES, but not physically identical IS, enhances dominance one week later, with PL activation being required during ES.

In contrast to males, prior ES in females had no impact on resistance behavior and spot occupancy one week later. Rather, all female groups (ES, IS, HC) displayed similar levels of dominance. This was not due to sex differences in learning the wheel-turn controlling response. Females readily acquired and maintained the instrumental controlling response to the same extent as males (Fig. 2*B*). However, prior work supports the idea that the acquisition and operation of control in females (i) does not lead to the same stress-buffering effects as in males and (ii) is accomplished with a different instrumental learning system than males, namely a dorsolateral striatum (DLS) ‘habit’ system rather than a corticostriatal ‘goal-directed’ system (Fallon et al., 2020; McNulty et al., 2022). If the DLS is inactivated, shifting instrumental performance to the DMS, control then leads to protection in females. Although speculative, future work should investigate whether ES in females would facilitate later dominance if corticostriatal structures (PL and DMS) were activated during the ES experience.

As mentioned, there is similarity in the mechanisms that mediate winning in social encounters and the protective effects of behavioral control over nonsocial aversive events. This led us to hypothesize that corticostriatal structures (PL and DMS) would be required for the development of stable dominance. We show that for winners of the initial warm spot competition, subsequent inactivation of the PL or blockade of the NMDA-dependent glutamatergic signaling in the DMS, the downstream striatal target of the PL, led to a lowering of dominance rank. Reduced status extended to subsequent drug-free competitions, thus interfering with the stability of dominance observed in vehicle-treated winners. The PL and DMS are known to mediate the effects of behavioral control because they mediate goal-directed learning, the form of learning in which instrumental effort is determined by the outcome. The PL and DMS doubtlessly participate in other processes, and it could be these that are important for winning in social encounters. However, dominance interactions are learning experiences as well as social interactions, and perhaps disruption of corticostriatal function interferes with the subject’s ability to use information from their previous success – such as the contingency between instrumental effort and outcome (e.g., occupying the warm spot) – to guide their performance in subsequent competition. The role of the corticostriatal system in aversively, rather than appetitively, motivated tasks is not well studied. However, the same manipulations used here that interfered with repeated winning also eliminate the resilience-producing effects of behavioral control over stress (Amat et al., 2005, 2014).

The above data suggest that repeated winning in the warm spot test might mimic the protective effects of behavioral control. It is known that reduced social investigation produced by IS is mediated by potent activation of 5-HT neurons in the DRN (Amat et al., 1998; Christianson et al., 2008, 2010), and this activation is inhibited by control. Indeed, we found that a history of winning prior to IS exposure blocked the increased levels of extracellular 5-HT in the DRN produced by IS and prevented social avoidance. These buffering effects were specific to winning, as repeated losing produced a very different neurochemical and behavioral pattern. Losing, independent of IS exposure, led to reduced juvenile social exploration, a finding that replicates previous work in mice (Šabanović et al., 2020). Losing also failed to modify the DRN 5-HT response to IS, with elevated 5-HT levels persisting throughout the entire shock session. It should be noted that other acute stressors, such as social defeat, activate DRN 5-HT cells (Cooper et al., 2009; Amat et al., 2010; Paul et al., 2011) and produce similar behavioral outcomes as does IS such as social avoidance and shuttlebox escape failure (Amat et al., 2010). This defeat-induced DRN 5-HT activation is selectively suppressed in animals with dominant status (Cooper et al., 2017). Taken together, these findings highlight several properties of winning as a resilience factor. Competitive success buffers against adversity that takes place in a novel context (generalization) and provides resistance to the effects of both nonsocial and social adverse events.

The impact of winning on the stressor response was selective. Neuroendocrine and central immune responses are also elicited by adverse events, although changes to these measures are typically not sensitive to the controllability of the stressor (Maier et al., 1986; Helmreich et al., 1999; Frank et al., 2007). Different energetic requirements are associated with social status and competition outcomes, and the levels of glucocorticoids and innate immune function can play a correlative or even causal role in the formation of hierarchies (Wingfield et al., 1990; Avitsur et al., 2007; Audet et al., 2010; Knight and Mehta, 2017). We assessed whether dominance status would alter IS-induced changes to circulating corticosterone and hypothalamic cytokine (IL-1β, IL-6) and chemokine (CX3CR1) mRNA expression levels. All measures (except IL-6) were impacted by IS, but the magnitude of change did not differ between winners and losers. The foregoing suggests that dominance status does not make the IS experience more or less “aversive” or “stressful”; rather it constrains the circuit (serotonergic) response to IS, thereby preventing the behavioral outcome (social avoidance). There are any number of experience-dependent intrinsic and extrinsic mechanisms that would alter how the dorsal raphe responds to IS. One possibility is that repeated winning experiences might strengthen PL top-down regulation of the DRN during IS (Grizzell et al., 2020), just as does prior ES.

In summary, our findings demonstrate that winning in social encounters and control over nonsocial stressors are interchangeable. The operation of behavioral control over adverse events, a key aspect of coping, facilitates later competitive success. Repeated winning in social encounters produces resilience against the nonsocial stressors such as uncontrollable shock. Given that control and stable dominance are implicated in positive health outcomes, the involvement of corticostriatal structures for both may represent a circuit-level endophenotype in the production of resilience.

## Acknowledgments and support

This work was supported by National Institutes of Health grants R01 MH050479 (SFM) and R21 MH116353 (MVB). The authors would like to thank Bryce Lorenz for technical support.

